# Fungal taste for minerals: the ectomycorrhizal fungus *Paxillus involutus* triggers specific genes when extracting potassium from different silicates

**DOI:** 10.1101/2021.03.05.434133

**Authors:** F. Pinzari, A.D. Jungblut, J. Cuadros

## Abstract

Silicates make up about 90% of the Earth’s crust and constitute the main source of mineral nutrients for microorganisms and plants. Fungi can actively weather silicates to extract nutrients. However, it is unclear whether they are able to obtain the same amounts of nutrients and use the same mechanisms when tapping into different mineral sources. We performed a microcosm experiment using the ectomycorrhizal basidiomycetes *Paxillus involutus* and the silicates K-vermiculite, muscovite and phlogopite as only potassium sources, as they show a different resistance for the removal of K cations from the mineral structure. A combination of transcriptomic, elemental and SEM analyses showed that different minerals stimulated specific weathering mechanisms and led to a change in fungal genes expression. The differential expression of the fungal genes generated alternative chemical attacks on the minerals, resulting in a tailored dissolution and selective uptake of chemical elements according to the leachability of K from the silicate mineral. The K uptake capacity of the fungus was highest with vermiculite in comparison to growth on phlogopite and muscovite. The findings provide new insights into fungal-mineral interactions that will help to interpret key processes for the homeostasis of soil environments.

## 1. Introduction

Weathering of silicates by fungi is an important process in soil formation and chemical evolution of rocks [1–3]. The ability to dissolve minerals applies to many fungi and in particular to mycorrhizal fungi, which form mutual symbiotic associations with the plant root system and are able to increase the dissolution and transformation of silicate minerals while extracting phosphorus (P), potassium (K), calcium (Ca), magnesium (Mg) and iron (Fe), especially under nutrient limitations [2]. The study of the genomes of ectomycorrhizal fungi and their transcripts have shown how fungi evolved complex mechanisms for the assimilation and exchange of carbon (C), P and nitrogen-based compounds [4]. However, metal nutrients have received much less attention, and their biogeochemical routes are almost unknown [2]. Free-living and mycorrhizal fungi can use a range of mineral dissolution mechanisms, including acidification of the microenvironment via excretion of protons, phosphoric acid, organic acids, and carbon dioxide (CO2) [5–7]. Fungi also use the production of extracellular polymeric substances to adsorb and accumulate cations and decrease the saturation for those elements [8]. Exudation of organic complex-forming molecules [8–9] is as well an essential mechanism for the modification of water chemistry (e.g., concentrating salts) and viscosity within biofilms that increases water reactivity with the mineral surface [10]. In addition to multiple mechanisms, there also seem to be different intensities of mineral weathering by fungi [11]. The production of oxalic acid by a fungus, for example, is modulated according to the type of mineral it comes into contact [12]. It is also known that there are genes that can be associated with specific weathering mechanisms and that these can be regulated by the depletion of certain nutrients from the culturing media [13]. What is unclear is whether, in light of the different structural ways in which nutrients may be part of the mineralogy of silicates, the fungi not only are able to modulate the intensity of weathering but also to implement different functional strategies. The ability of fungi to change the metabolic strategy depending on the type of mineral from which they derive nutrients would have implications for both the ecology and evolution of fungi and fungi-plant systems [6, 14]. Fungal-derived chemical equilibria in the biosphere would vary based on gene expression mechanisms and regulatory processes whose control would disclose new frontiers in agriculture and soil management. In fact, a better understanding of the regulation of weathering processes by fungi could improve predictions of ecosystem functions [15–16]. An active fungal regulation in silicates weathering would also impact Earth system models and provide new variables to the global C cycle [17–19].

This study tested the hypothesis that fungi would show distinct mechanisms of attack and differential gene expression profiles when grown on specific minerals to obtain limiting nutrients such as K. The fungus *Paxillus involutus* was selected for the experiments as it is a Basidiomycetes of global significance [20]. It can form mycorrhizae with many species of trees [21] but is capable also of growing in axenic culture [22] and can weather minerals [12, 23–25]. *P. involutus* was grown in microcosm experiments under three different conditions where the only sources of K were the minerals phlogopite, K-vermiculite and muscovite, and compared with a positive control with K available in the media and a negative control without any K. These silicates have similar metal nutrient contents but have different degrees of weatherability, which could trigger a specific fungal strategy to obtain nutrients and reveal a differential genetic expression. The growth and response of the fungi were evaluated using RNAseq-based differential gene expression. The mycelium was analysed to evaluate the uptake of nutrients using inductively coupled plasma mass spectroscopy (ICP-MS), and the minerals were analysed to detect traces of weathering and patterns of interaction with the fungal hyphae by means of scanning electron microscopy (SEM).

## 2. Materials and Methods

### 2.1 Fungal material and experimental set-up

Cultures of *P. involutus* (Batsch) Fr. (strain ATCC 200175, www.atcc.org) (Basidiomycota, Boletales) were reactivated and maintained aseptically on Modified Melin–Norkrans (MMN) agar (Supplementary Materials S1). The purity and identity of *P.involutus* ATCC 200175 strain was verified by Sanger sequencing (Supplementary Materials S2). The microcosms were prepared according to Wei et al. [26] with some modifications (Supplementary Materials S3). Three different minerals with decreasing level of K leachability were used: muscovite, phlogopite, K-vermiculite. Their structural formulas are specified in Supplementary Materials S4. All cultures were kept in the dark at 25°C throughout the experiment. The microcosms were incubated for 21 days. Ten biological replicates from each experiment were used for RNA extraction, five from each experiment were dismounted and used for ICP-MS analysis, and the corresponding minerals were used in SEM analysis. After the experiments, fungal biomass was determined by weight using five microcosms from each experiment (dry weight, 50°C, 48 h). The pH of the agar was measured at the beginning and the end of the experiments (Electrode n.662-1164, VWR International Ltd, Leicestershire, U.K.).

### 2.2 Microscopic and chemical analysis of fungal biomass and minerals

Mineral samples were analysed before and after the experiment using SEM as described in Supplementary Materials S5. Groups of five replicates from each experiment were used to analyse sodium (Na), Mg, aluminium (Al), Fe, P, K, and Ca in the fungal mycelium by digestion and analysis by ICP-MS (Agilent Technologies 7700), as described in Supplementary Materials S6.

### 2.3 RNA extraction and libraries preparation

At the end of the experiment, fungal biomass was removed from the cellophane membrane and ground into a powder in liquid nitrogen. mRNA was extracted using the RNAeasy^®^ PlantMini Kit (Qiagen, Manchester, U.K.) and libraries prepared using TruSeq Stranded mRNA Kit (Illumina, Cambridge, UK) (Supplementary Materials S7 and 8). Libraries (5 biological replicates for each experiment, for a total of 25 samples) were sequenced with Illumina NovaSeq 6000 (2 x 100 bp readlength) by the CeGAT Company (Germany).

### 2.4 Bioinformatic analysis

Full details on the sequence processing are provided in Supplementary Materials S9. Trimmed reads were aligned to the *Paxillus involutus* reference genome (GCA_000827475.1) using STAR (v2.5.2b) [27]. A count table was created using the HTSeq package [28] implemented in OmicBox (version 1.2.4) [29–30]. EdgeR version 2.12 [ref. 31] was used to perform pairwise differential expression analysis (DEA) between pairs of experimental conditions (5 replicates per experimental condition). Functional annotation of differentially expressed genes (DEGs) was based on *P.involutus* annotation from the EnsemblFungi server (http://fungi.ensembl.org) with assembly accession number GCA_000827475.1 [ref. 21], and by using the BLASTx algorithm within OmicsBox version 1.2.4 (BioBam) [29–30]. Functional annotation of sequences was further achieved using InterProScan [29–30, 31] and EggNOG-mapper [32]. Gene Set Enrichment Analysis (GSEA) was used to determine whether an *a priori* defined set of genes showed statistically significant, concordant differences between two biological states [34]. The normalised enrichment score (NES) was used to compare analysis results across gene sets. The false discovery rate (FDR), was used as an estimate of the probability that a gene set with a given NES represented a false positive finding. FDR cut-off was fixed at 5%.

## 3. Results

### 3.1 Fungal growth

*P. involutus* mycelium grew in all experiments showing good replicability of the in-vitro system utilised (Fig. S1). The pH of the agar decreased in all the experiments from its initial value of 4.7. Table S1 shows the pH values in the five experiments at the end of the incubation, with a higher pH in the positive control (Cp) than in the microcosms with the minerals. The pH values in the treatments with minerals were significantly different from the positive control, whereas the pH values in the negative control (Cm) were not significantly different from the positive control or the experiments with the minerals. The comparison of the fungal biomass content obtained in the five experiments is shown in Table S2. The wet fungal biomass was significantly higher in the positive control, followed by the experimental treatment with phlogopite, while significantly lower values were measured in the K-vermiculite, muscovite and negative control experiments.

### 3.2 Fungal weathering of minerals: ICP-MS results

The fungus absorbed more K from K-vermiculite than from the other two minerals (Table 1). There was a trend indicating a slightly higher concentration of K in experiments run with phlogopite than muscovite (Table 1), but the K content of mycelium grown in the presence of muscovite and phlogopite was not significantly different from that recorded in the negative control. Table 2 shows the Pearson correlation coefficients (*r*) calculated between the mycelial concentrations of the elements. The coefficient values showed that the absorption of K was positively correlated with P, Ca and Mg, while it was negatively correlated with the absorption of Na. Moreover, Fe and Al showed a strong positive correlation (r=0.99).

**Table 1.** Uptake of elements by the fungal mycelium. Values are the means (±SD) from n = 5 replicates. Means in a column without a common letter (a/b/c) differ for P < 0.05, as analysed by one-way ANOVA and Tukey’s post hoc test.

|  | K | Mg | Al | P | Na | Ca | Fe |
| --- | --- | --- | --- | --- | --- | --- | --- |
| *ppm ($\mu\text{g g}^{-1}$ of dry mycelium)* |  |  |  |  |  |  |  |
| Cp | 304.11±11.05 a | 7.16±0.35 a | 0.06±0.001 b | 110.54± 5.70 a | 6.55±0.38 c | 3.20±0.17 a | 0.33± 0.04 b |
| Cm | 5.52±1.69 c | 5.82±0.75 a | 0.07±0.01 b | 70.35± 10.49 b | 47.76±11.47 b | 2.99± 0.52 ab | 0.33± 0.04 b |
| M | 6.80±0.96 c | 6.21±0.85 a | 0.28±0.09 b | 73.24± 11.07 b | 51.35±8.31 b | 2.56± 0.41 b | 0.48± 0.09 b |
| P | 8.64±2.91 c | 6.67±1.15 a | 1.12±0.78 a | 68.57± 7.90 b | 47.64±8.00 b | 2.73± 0.23 ab | 0.93± 0.50 a |
| V | 25.11±4.31 b | 6.75±0.47 a | 0.16±0.04 b | 79.52± 4.52 b | 64.57±4.81 a | 2.63± 0.23 ab | 0.41± 0.06 b |
| Pr > F (Model) | < 0.0001 | 0.10 | 0.0001 | < 0.0001 | < 0.0001 | 0.04 | 0.0001 |
| Significant | Yes | No | Yes | Yes | Yes | Yes | Yes |

**Table 2.** Correlation coefficient (Pearson) between elements concentration in the fungal mycelium across all the experiments. Values range from −1 (complete negative linear correlation between variables) to 1 (complete positive linear correlation).

| Variables | Na | Mg | Al | P | K | Ca | Fe |
| --- | --- | --- | --- | --- | --- | --- | --- |
| Na | **1** | -0.08 | 0.20 | **-0.59** | **-0.87** | -0.33 | 0.22 |
| Mg | -0.08 | **1** | 0.39 | **0.67** | **0.42** | **0.54** | **0.43** |
| Al | 0.20 | 0.39 | **1** | -0.28 | -0.28 | -0.10 | **0.99** |
| P | **-0.59** | **0.67** | -0.28 | **1** | **0.89** | **0.63** | -0.26 |
| K | **-0.87** | **0.42** | -0.28 | **0.89** | **1** | **0.49** | -0.28 |
| Ca | -0.33 | **0.54** | -0.10 | **0.63** | **0.49** | **1** | -0.10 |
| Fe | 0.22 | **0.43** | **0.99** | -0.26 | -0.28 | -0.10 | **1** |

*Values in bold are statistically significant ( $p < 0.05$ )*

### 3.3 Scanning Electron Microscopy of mineral surfaces

The SEM imaging showed that the three minerals were altered by the growth of the fungus during the microcosm experiments. Comparing SEM images of minerals before and after the experiments suggested that *P. involutus* when forced to grow under limited availability of K, produced different effects in each silicate mineral. In muscovite, the surface of the crystal appeared eroded and uneven (Fig. 1A). At higher magnification, alterations were observed at the contours of the fungal hyphae consisting in the localised removal of surface layers of the mineral (Fig. 1B). In phlogopite, large portions of mineral layers were removed (Fig. 1C). In the high-resolution images, the surface of the phlogopite crystals was fragmented and furrowed, and characteristic depositions arranged in small regular clusters were observed between the cracks (Fig. 1D). Fungal mycelium appeared singularly adherent to the crystal when grown on the K-vermiculite. Here, forms of surface erosion were also visible at low magnification suggesting that the fungus exerted a mechanical action (Fig. 1E). However, the waxy consistency of the superficial layer of K-vermiculite, at times excavated and raised to form small accumulations of material, suggests that a chemical action also took place to make an initially intact crystalline structure malleable. High magnification images of K-vermiculite surface show some hyphae flattened and others filled with precipitated material (Fig. 1F).

**Figure 1.**
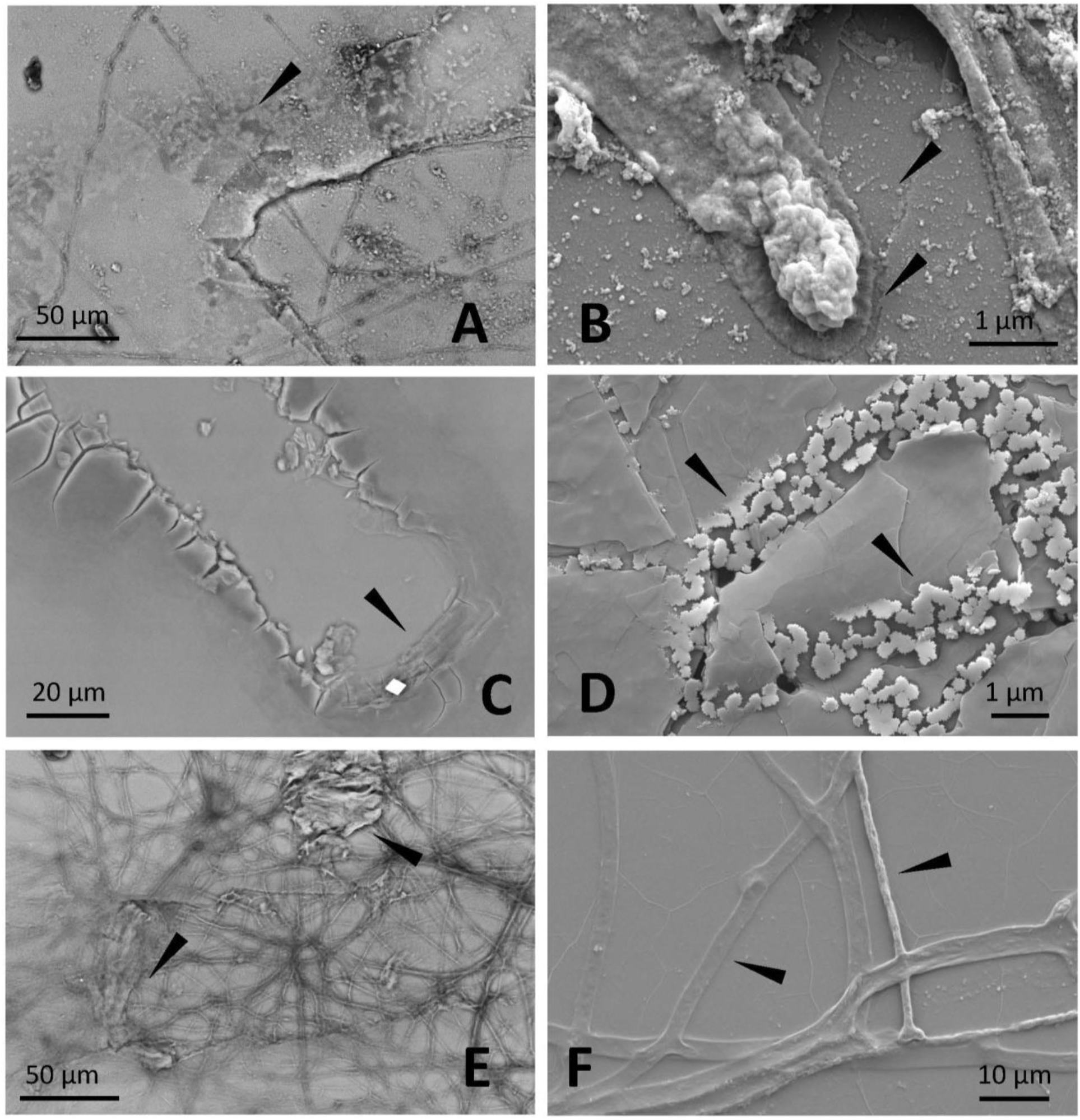
Scanning Electron Microscopy imaging of the minerals after fungal growth. On the left (A, C, E), images obtained in variable pressure mode with backscattering detector, on non-coated samples. On the right (B, D, F), images captured in a high vacuum on gold-palladium coated samples. A) muscovite: the arrow indicates superficial erosion of the mineral that occurred after the exposure to fungal activity; B) muscovite: the arrows point to the tip of a fungal hypha attached to the surface of the mineral, and the superficial trenching; C) phlogopite: the arrow points to the edge of a furrow caused on phlogopite surface by the fungus; D) phlogopite: the arrows indicate regular clusters of depositions between the crack caused by the fungus onto the surface of the crystal; E) K-vermiculite: the arrows indicate surface erosion, with a localised scratching and slipping of the surface layer of the crystal; F) K-vermiculite: the arrows indicate fungal hyphae adhering to the mineral’s surface, with some flattened, also due to the high vacuum, and others filled with some precipitated material.

### 3.4 *Differential expression* of P. involutus *during K weathering on K-vermiculite, muscovite and phologpite minerals*

The transcriptome sequencing yielded 966.96 million reads, with a total number of bases (in Gb) of 194.89, an average GC content of trimmed FASTQ reads (average of all samples) of 53.0% to 54.0% and a Q30 quality score threshold value between 91.3% and 93.07% (Table S3). Multidimensional scaling analysis (MDS) (Fig. 2) [35] was performed to evaluate the dissimilarity between transcriptomes of *P.involutus* obtained in response to its growth on K-vermiculite, phlogopite and muscovite and in comparison to the negative and positive controls. In the two-dimensional scatterplot (Fig. 2) the distances represent the log2 fold change (log2 FC) between samples. The transcriptomes obtained with minerals were different from the positive control. The transcriptome from the K-vermiculite microcosm showed a high similarity between replicates and grouped separately from all the other experiments. The transcriptomes from phlogopite, muscovite and the negative control overlapped and showed a general higher variability compared to positive control and K-vermiculite treatment.

**Figure 2.**
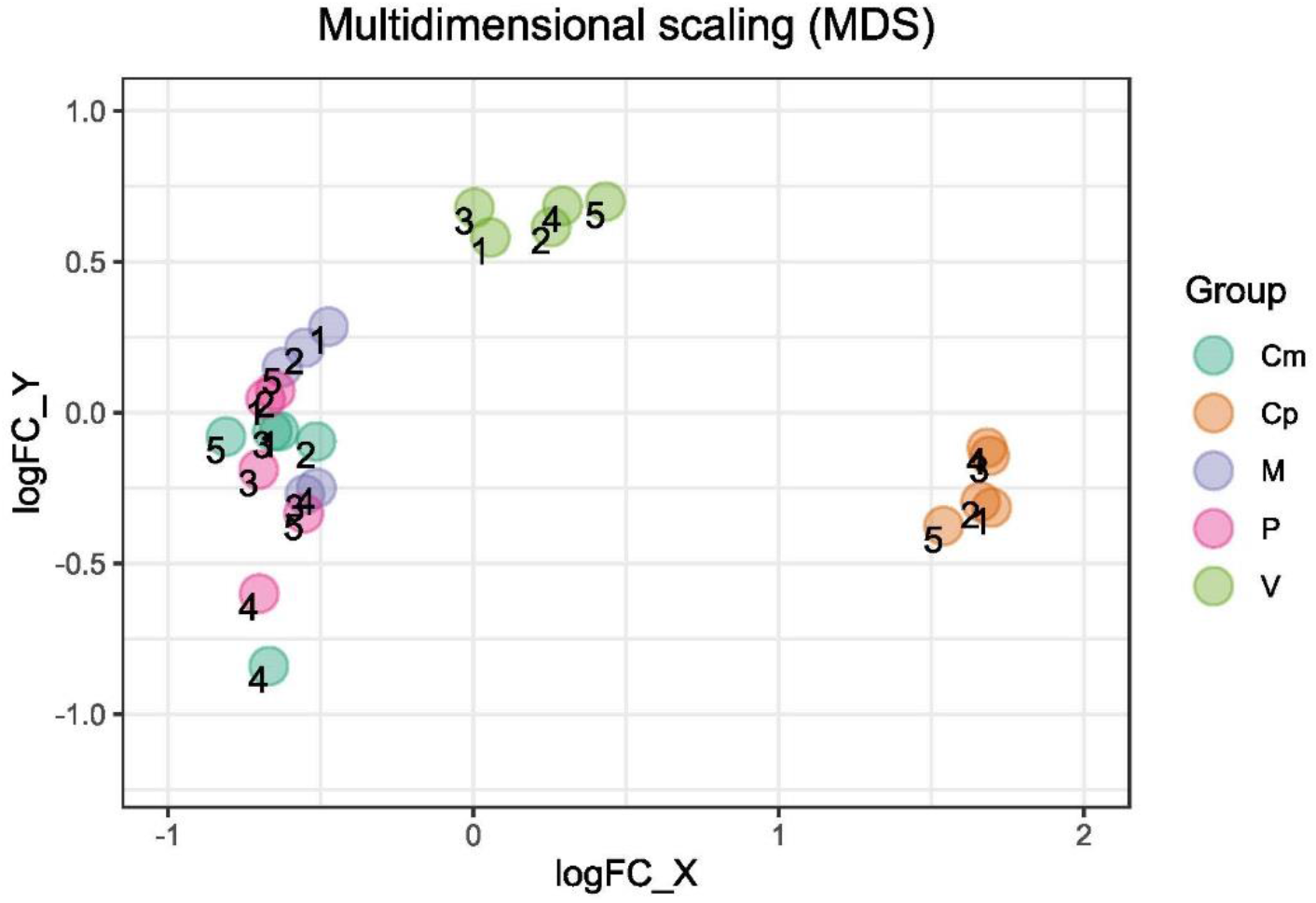
Multidimensional Scaling analysis of transcriptomes obtained from the *P. involutus* grown K-vermiculite, muscovite and phologpite as well as positive and negative controls. The distances represent the log2 fold change between samples. The symbols differentiate between experiments, the numbers represent the biological replicates within each experiment.

From the heatmap (Fig. 3) emerges that there are two significant clusters of genes with distinct regulation for the positive and negative controls. The fungus in the presence of K-vermiculite showed a gene expression partially similar to the positive control. With muscovite and phlogopite the gene expression was similar to the negative control. The number of genes differentially expressed by the fungus in each comparison are shown in Table 3. Overall, the number of genes differentially expressed between the experiments with the minerals and the negative control is lower than the number of genes obtained in the contrasts against the positive control. The similarity between the condition of K deprivation and that of bioavailability conditioned by the mineral source is also shown by the Volcano plots that display both the genes differentially expressed (FDR < 0.05) and those that did not show a change in their expression level between contrasting conditions (Figure S2). The number of significantly up and down-regulated genes (FDR < 0.05) between the experiments and controls is summarised in Table 3. In the pairwise comparison between transcriptomes of *P. involutus* grown on the three minerals, and the negative control, the number of differentially expressed genes decreases from K-vermiculite (439) to phlogopite (47) and muscovite (29). There were very few DEGs between muscovite and negative control, which indicates that the physiological state of the fungus in the presence of muscovite was similar to being under the condition of K deprivation.

**Figure 3.**
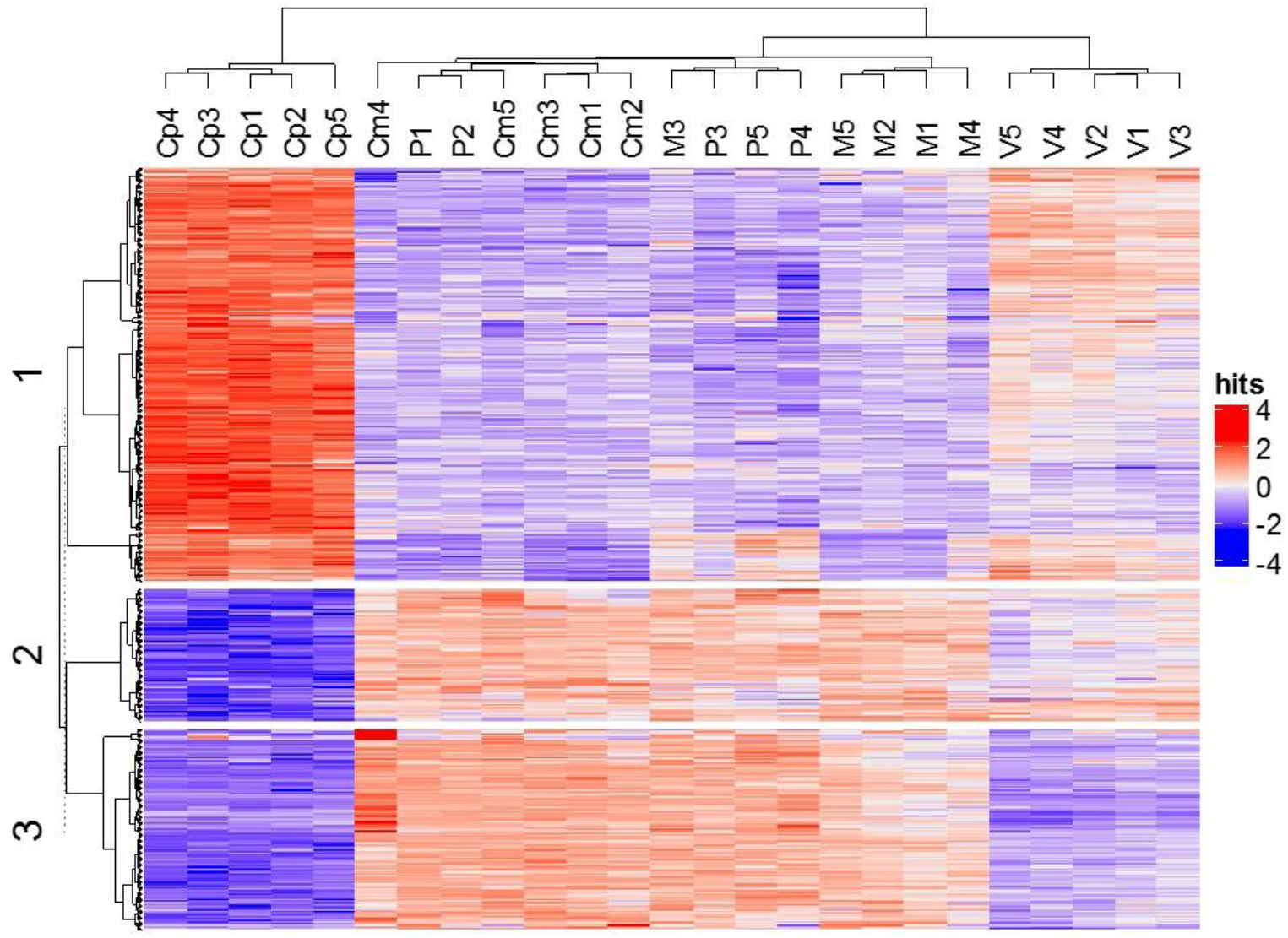
Heatmap of the top 1000 differentially expressed genes (ranked by FDR). It is a twodimensional visual representation of data in which a range of colours represents numerical values of points. The dendrograms added to the left, and top are produced by a hierarchical clustering method based on Euclidean distance computed between the differentially expressed genes (left) and samples (top).

**Table 3.** Number of up- and down-regulated genes between pairs of treatments (FDR < 0.05), positive control (C plus K= Cp), K-vermiculite (V), phlogopite (P), muscovite (M) and negative control (C minus K= Cm). In parenthesis those genes that were up- and down-regulated for more than 2 logFC, without parenthesis those genes that were up- and down-regulated for more than 1 logFC. Total DEGs is the sum of up and down regulated genes for (FDR < 0.05).

|  | Up-regulated | Down-regulated | Total significant DEGs | Total genes shared between pairs of conditions |
| --- | --- | --- | --- | --- |
|  | Log FC>1<br>(Log FC>2) | Log FC>-1<br>(Log FC>-2) |  |  |
| M/Cm | 21 (0) | 8 (5) | 29 | 968 |
| P/Cm | 39 (1) | 8 (5) | 47 | 1335 |
| V/Cm | 191 (11) | 248 (35) | 439 | 5008 |
| Cp/Cm | 832 (111) | 676 (155) | 1508 | 7536 |
| M/Cp | 583 (103) | 744 (120) | 1327 | 7355 |
| P/Cp | 730 (103) | 891 (120) | 1621 | 7732 |
| V/Cp | 198 (39) | 428 (62) | 626 | 5786 |

Fig. 4 and Supplementary Spreadsheet 1 show the genes that the fungus either triggered or repressed at the presence of the three minerals and the positive control, compared to the negative control based on FDR values and log2FC scores. The differential expression between positive and negative control identified marked differences between the condition of total bioavailability of K and its unavailability, with genes showing up to 6 log2FC values. However, the top genes were assigned as hypothetical proteins without associated specific function.

**Figure 4.**
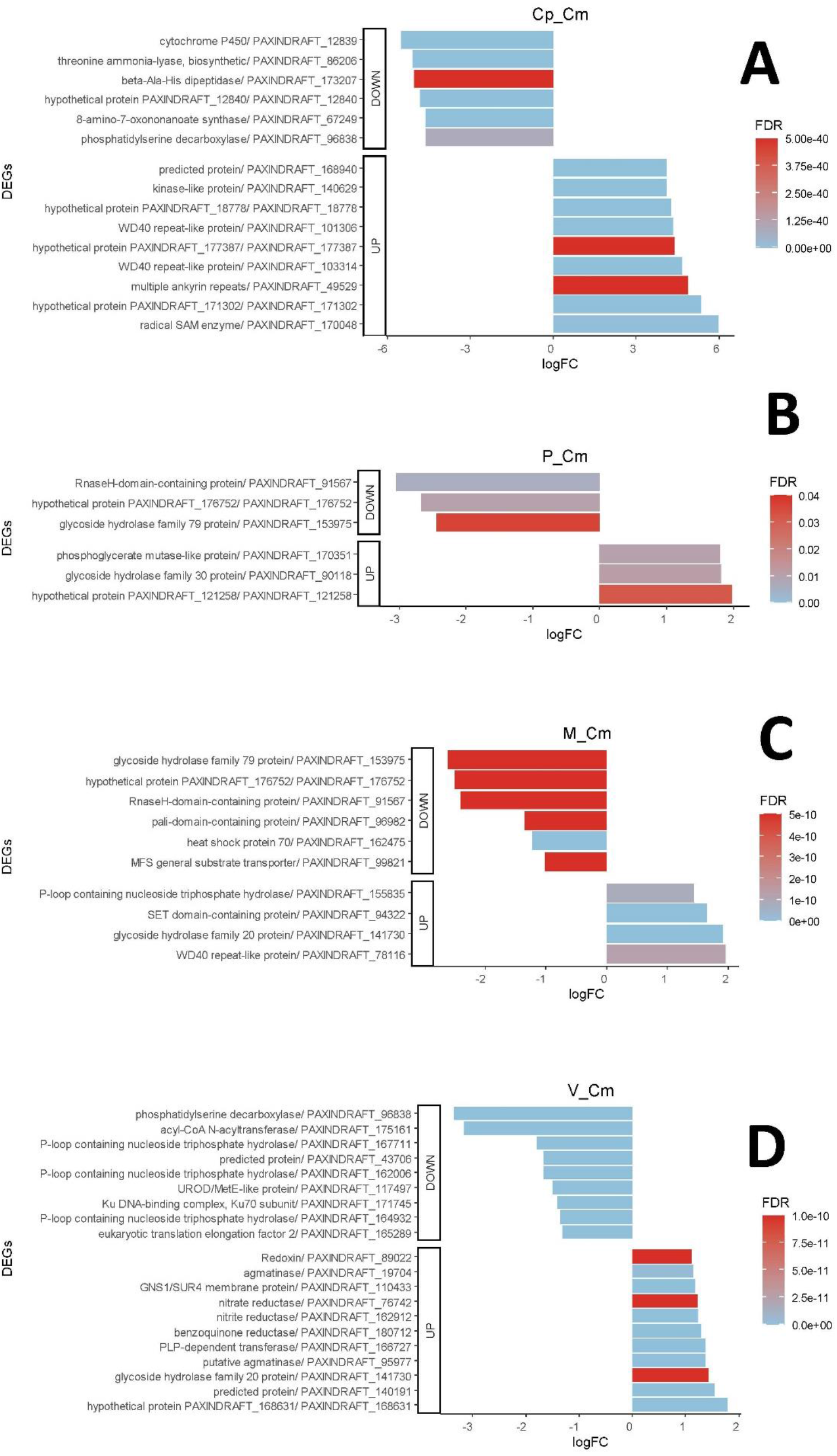
Top Up- and Down-regulated genes between the experiments in which the fungus grew in the presence of the three minerals (V, P and M) compared to the negative control situation (Cm).

### 3.5 Gene set enrichment analysis (GSEA)

The genes that were identified as differentially expressed between pairs of treatments were grouped by their involvement in biological processes, molecular functions and cellular compartments. In Fig. 5 and Supplementary Spreadsheet 2 are shown the normalised enrichment score values (NES) obtained for comparisons between the experiments in which the fungus grew in the presence of the three minerals (V, P and M) compared to the negative control condition (Cm). Among the molecular functions stimulated in the experiments with phlogopite there were carboxylic-acid metabolic process (GO:0019752), organic-acid metabolic process (GO:0006082), oxoacid metabolic process (GO:0043436) and lipid metabolic process (GO:0006629). The molecular functions stimulated in the case of muscovite were different and significantly fewer, compared to the other two minerals. The only molecular functions overexpressed in *P.involutus* when incubated with muscovite were monooxygenase activity (GO:0004497) and oxidoreductase activity (GO:0016705-MF). Interestingly, the biological processes that were under-expressed (compared to the negative control condition) were response to heat (GO:0009408), cellular response to heat (GO:0034605) and ribonucleoprotein complex subunit organisation (GO:0071826). There were also differences observed in the gene assignment for the fungus grown with K-vermiculite as the only source of K. In this condition, both molecular functions and biological processes were associated with the cytoskeleton, and significant overexpression of gene sets related to cytoskeletal protein binding (GO:0008092) and cytoskeleton organisation (GO:0007010). Moreover, with K-vermiculite, catabolic process (GO:0009056), vesicle-mediated transport (GO:0016192), and cellular response to stimulus (GO:0051716 i.e. processes resulting in a change in state or activity of a cell) were also overexpressed.

**Figure 5.**
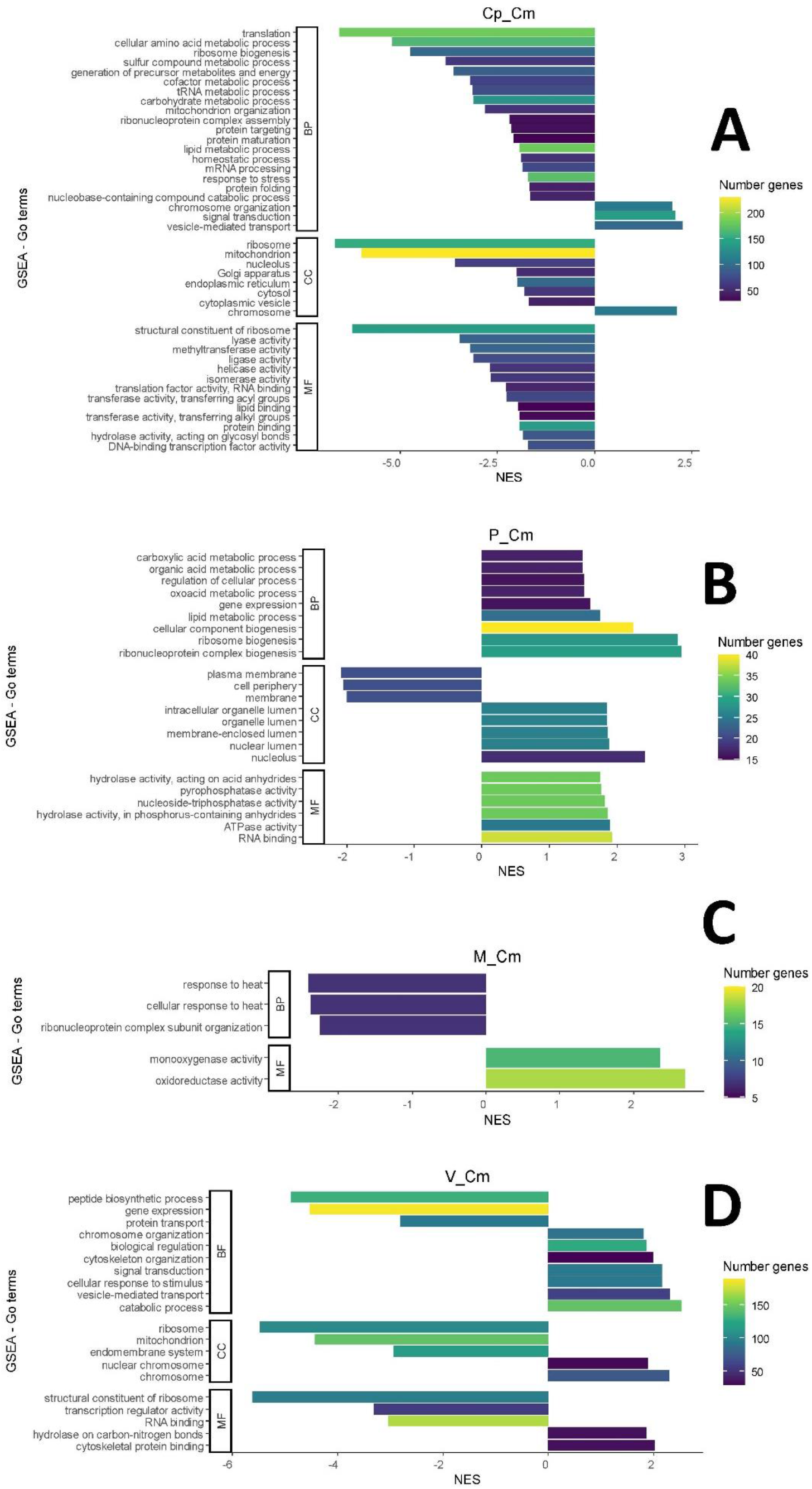
Gene Set Enrichment Analysis (GSEA). Treatments with the three minerals (V, P and M) and the positive (Cp) control, compared to the negative (Cm) control condition.

## 4. Discussion

The present study confirmed that *Paxillus involutus* is able to weather K from silicates such as K-vermiculite, muscovite, and phlogopite, but we also showed for the first time that the gene expression patterns and metabolic response differed according to K leachability from different types of silicate minerals. Differences in chemical and physical attack by the fungus on the three minerals were also documented by chemical and imaging analyses.

### 4.1 Effects on pH and uptake of K and other mineral nutrients

The pH of the agar under the mycelium decreased significantly in all K-depletion treatments, which is likely due to secretion of acids by the fungus in response to either the absence of K or the presence of K containing minerals in the agar. This stimulation of acid secretion appeared stronger in the presence of minerals than in the negative control, which reinforces the idea that it is not the absence of K, but the presence of minerals containing it, that stimulated the secretion of acids into the culture medium. K concentration in the mycelium was negatively correlated with Na concentration, but positively correlated with P. There was no clear correlation observed between Mg and Ca. The negative correlation of K with Na was likely due to the use of Na as a substitute for K by the fungus. This can be explained by K being the primary inorganic cation in the fungal cell cytoplasm and an essential macro-element that is accumulated against a transmembrane concentration gradient. If the K deficiency occurs in the presence of Na, fungal cells can take Na+ instead of H+ as a substitute for the missing K [36]. While fungi can utilise Na to grow in the absence of K, Na is also a toxic element in excessive quantities [37]. Thus, the recorded changes in gene expression that depended on K limitation could have been linked to functional modifications needed at membrane level to assimilate Na instead of K, and mechanisms to react to Na toxicity [38].

We also observed a positive correlation of K and P, which could be due either to differential P uptake, or to differential P excretion. The findings agree with previous studies, where a strong correlation between P and K cellular levels was observed for the ectomycorrhizal fungus *Hebeloma cylindrosporum* [39]. Jentschke et al. [40] found that the fungus *P. involutus* translocated K and Mg to Norway spruce plants, with the translocation of these elements depending on their coupling with P uptake during the process. Our findings also suggest that K was one of the major counter-ions of polyphosphate (polyP) granules, especially of soluble polyP short-chains, mainly located in fungal vacuoles [41]. K and P have been shown to be located in the same vacuolar fungal compartments of *P. involutus* [42–43]. P uptake and polyphosphates synthesis and accumulation could result in the accumulation of a high negative charge in the cell, which results in the activation of cation import mechanism for maintaining overall cellular charge neutrality. Therefore, K is both an element directly involved in fungal nutrition, but also a component necessary for cell homeostasis and efficient uptake and storage of inorganic phosphate. Although the fungus grew in K-depleted experiments and P was not limiting, less P was assimilated than in the positive control. This indicates that the limitation in the bioavailability of K could have affected P assimilation.

The SEM imaging allowed to document visible effects of metabolic action by the fungus on minerals at microscale level. The K extraction led to the formation of furrows, cracks, swelling, erosions and bio-precipitation of material on crystals. Some of the effects are due to the production of organic acids, the adhesion of hyphae and leaching of elements other than K. For example, the mycelium grown in the presence of phlogopite showed significantly higher assimilation of Fe and Al than the other experiments. Ectomycorrhizal fungi can assimilate Fe [44] and are also capable of absorbing Al [45]. Similarly, Bonneville et al. [44] showed that *Paxillus involutus* could oxidise Fe(II) to Fe(III) in biotite, increasing with this process the mineral weathering. The authors suggested that Fe(II) oxidation induces the loss of the newly formed Fe(III) from the octahedral sheets and formation of amorphous Fe oxy-hydroxides within the silicate layers, which induces volume increase, strain in the biotite lattice and the formation of fractures.

### 4.2 *Differential gene expression of* P. involutus *as a response to growth on silicates with variable K leachability*

The gene expression profiles by *P. involutus* were distinct for mycelium grown on K-vermiculite in comparison to muscovite and phlogopite as well as to controls. The muscovite and phlogopite experiments had similar profiles and overlapped with the negative control. The differential expression of genes matched to the concentration of K found in the mycelium after the experiments, and there was a positive relationship between the accessibility of the element and its absorption by the fungus. This suggested the fungus modulated the gene expression according to both the amount of K that it was able to assimilate from the silicates but also reflected the strength with which the three minerals retained the cation. As the three minerals showed some variation in the weathering of other elements such as Fe, Mg, Na and Al, the gene expression profile might also be affected by other factors than leachability of K from the respective mineral, including toxicity of released metals, and solubility of the other cations in the growth environment [46–47].

In the presence of K-vermiculite, the fungus was able to derive the K by its displacement from the interlayer sites of vermiculite by other cations in the medium, which is an exchenge mechanism similar to how fungi obtain minerals in soil environments [48]. This suggests that the fungus was not subjected to the same level of K deprivation as with the other two minerals. However, it still had to activate different mechanisms for the assimilation of K than under condition of its unlimited bioavailability (in positive control). In the case of muscovite, which is the mineral most strongly retaining K, the gene expression profiles suggested a response similar to growth in the absence of K. There were only 21 genes up- and 8 down-regulated by the fungus in the presence muscovite in comparison to the negative control, moreover, only a few of these were found to be differentially expressed for more than 2 logFC. In terms of K effectively being assimilated by the fungus, the difference corresponds to about one microgram per dry gram of mycelium. These few genes are therefore those that separate the condition of total K depletion from that of a minimum bioavailability. The most differentially expressed genes (downregulated more than 2 logFC in the presence of muscovite, compared to negative control) were a glycoside hydrolase family 79 protein from the membrane (PAXINDRAFT_153975, GO:0016020), a hypothetical protein associated to the nucleus, involved in regulation of transcription by RNA polymerase II and with DNA-binding transcription factor activity (PAXINDRAFT_176752, P:GO:0006357; F:GO:0000981; F:GO:0008270) and an RnaseH-domain-containing protein, with RNA phosphodiester bond hydrolytic function, binding nucleic acids (PAXINDRAFT_91567, P:GO:0090502; F:GO:0003676; F:GO:0004523). The glycoside hydrolase family 79 protein includes endo-beta-N-glucuronidase (EC 3.2.1.31) and enzymes with β-glucanase activity, that in fungi are thought to be necessary in morphogenetic events that require controlled hydrolysis of the cell wall [49].

Comparison of positive and negative control conditions showed that more genes were up regulated under unlimited K bioavailability than under stress conditions of K depletion. Which corresponds to a large down-regulation of genes in the negative control. This can be read as a response to stress with the deactivation of functions in order to overcome the nutrients limitation. Still, a significant down regulation of genes in the positive control microcosms experiments was also observed, indicating the existence of a number of mechanisms that have evolved to respond to the absence of K in the environment. However, the fungus still grew under K-depleted conditions as the dry weight of the mycelium in the negative control was not significantly different to biomass in the positive control. This could be due to the increased uptake of Na to replace K as cation, as Na content of the mycelium in the negative control was at least seven times that of the positive control. The change in gene expression triggered by K limitation could have therefore been linked to functional modifications needed at membrane level to assimilate Na instead of K, and to limit any toxic effects by the increased uptake of Na in to the cells. Similar mechanisms of reaction of *P. involutus* to limiting nutritional conditions have been observed by Paparokidou et al. [50] who showed that inorganic P starvation induces in the fungus expression of newly identified putative high-affinity Pi transporter genes, while reducing the expression of putative organic acid transporters.

### 4.3 Differences in functional genes and processes for K leaching from silicates

Phlogopite, K-vermiculite and muscovite led to distinct transcriptional responses in the fungus, suggesting some mechanism by the fungus to sense and respond to its mineral surroundings. Gene expression of the fungus in the presence of K-vermiculite compared to the condition of total K depletion highlighted the overexpression of genes that interact with cytoskeleton components which have a role in the assembly or disassembly of cytoskeletal structures such as actin. They are also involved in vesicle-mediated transport, and suggests that adhesion mechanisms could have had a role in K uptake. Indeed, fungi, in addition to dissolving the minerals with the production of acidic substances, have shown in some studies to carry out physical attacks on minerals through hyphal adhesion. It has been shown for example that the release of K was enhanced by a factor of 3-4 by direct contact between K-feldspar and illite surfaces and the fungus [51]. Attachment of fungi to the mineral surfaces has also been shown to cause a more efficient release of elements from biotite [52–53]. Physical mechanisms of action may be therefore particularly efficient for some cations such K.

Phlogopite triggered the overexpression of genes associated with the chemical reactions and pathways involving carboxylic acids, organic acids and also oxoacid metabolic process. An oxoacid is a compound which contains oxygen and which produces a conjugate base by loss of positive hydrogen ion(s). Many organic acids, like carboxylic acids, are oxoacids. Van Schöll et al. [25] showed how ectomycorrhizal fungi lacking mineral nutrients like P, Mg, and K might modify the composition of organic acids they exude. In other fungi, oxalate exudation was found to be higher when the mycelium was in contact with rock grains containing K [54]. Furthermore, the biological processes that were stimulated by phlogopite involved the hydrolytic activity linked to P-containing anhydrides for the catalytic reaction that produces nucleoside diphosphate and a phosphate from nucleoside triphosphate. There are the adenosine triphosphatases (ATPases) that generate electrical and chemical ion gradients across membranes and transporters for a wide range of solutes across membranes. Another gene up-regulated by the fungus in the presence of phlogopite, compared to the negative control, is a phosphoglycerate mutase-like protein, which was annotated as an integral component of the plasma membrane. The top up-regulated gene in K-vermiculite was also a hypothetical protein that is located in the membrane and is involved in ATP-coupled electron transport and electron transfer activity. This highlights the involvement in transport mechanisms during leaching and uptake of K from silicates.

Monooxygenase and oxidoreductase activities were the only molecular functions significantly overexpressed as gene sets in the presence of muscovite. Fungi produce heme-containing peroxidases and peroxygenases, flavin-containing oxidases, and different copper-containing oxidoreductases [55]. Moreover, in some basidiomycetes unspecific peroxygenases can catalyse a variety of monooxygenation reactions with H2O2 as the source of oxygen and final electron acceptor. Oxidation processes are more energy-demanding than acid production, and therefore they might have been used by the fungus after failure to leach K under acidic conditions, as suggested by the low uptake of K in the mycelium and the limited uptake of Al, compared to phlogopite, and by the evidence that fungal weathering of Al from muscovite occur by releasing organic acids [56]. Xiao et al. [57] and Wang et al. [58] suggested that genes encoding multicopper oxidase have a significant role in the weathering of K-bearing silicates in *Aspergillus* spp. Our study identified a gene encoding for di-copper centre-containing protein with oxidoreductase and metal ion binding function as up-regulated by the fungus grown with muscovite. Wang et al. [59] also showed with an electrochemical experiment that multicopper oxidases could enhance electrons transfer during fungal weathering of K-bearing silicates, promoting electron transport at the edge of the minerals.

There are four families of putative K transport systems identified in ectomycorrhizal fungi [60]. HAK and Trk transporters, TOK and SKC channels, the latter representing voltage-dependent K-selective channels. In particular, Trk transporters are membrane proteins involved in K uptake that are widespread among fungi [61]. HcTrk1, one member of these transporters that are present in some ectomycorrhizal fungi [62] was described as a Na+ /K+ transporter. However, in *P. involutus* none of the genes belonging to K transporters (HAK and Trk transporters, or to TOK and SKC channels) were significantly up-regulated in the microcosms with the minerals compared to the negative control. In contrast, several genes associated with transport systems were down-regulated in the mycelia grown with K-vermiculite compared to negative control, including a P-loop containing nucleoside triphosphate hydrolase protein associated to ATPase-coupled transmembrane transporter. There was also a gene down-regulated in the muscovite experiment that is associated with a general substrate transmembrane transporter (major facilitator superfamily MFS), and an integral component of the plasma membrane. The MFS are membrane proteins that import and export target substrates such as metabolites, oligosaccharides, amino acids and oxyanions. They utilise the electrochemical gradient of the target substrate (uniporter), or act as a co-transporter where transport is coupled to the movement of a second substrate.

The presented study is based on laboratory experiments, however the fungus *P. involutus* is abundant in the environment and forms mycorrhizae with many tree species. Many soil environments can be cation-poor and therefore the ability of *P. involutus* to access K from minerals would give an ecological advantage. Therefore, the fungi might have even evolved specific cellular and metabolic mechanisms [48, 63]. An adaptation of the fungi to low K conditions could help to explain the observed down regulation of many genes in the positive control microcosms experiments. The down regulated genes under unlimited K conditions were predicted to be linked to fundamental fungal biological processes, such as “translation”, “carbohydrate metabolic processes” or “ribosome biogenesis”. A large set of genes (over 200) associated to the “mitochondrion” cellular components were also significantly down-regulated. This suggest that the fungi might had a lower energy expenditure and cellular activity in the presence readily bioavailable K.

## 5. Conclusions

The geo-biological cycles are governed by complex interactions and feedback mechanisms between living organisms and rocks. Filamentous fungi play a key role as they collaborate symbiotically with plants, decompose and recycle organic matter and extract nutrients from minerals. One of the fundamental nutrients for growth is K, and the global transcriptome analysis showed that *Paxillus involutus* could switch on or off different genes and metabolic pathways depending on the minerals from which it is forced to obtain K. The differential expression of the fungal genes generated alternative chemical attacks on the minerals, resulting in a tailored dissolution and selective uptake of chemical elements. The fungus also showed an excellent exchange capacity with K-vermiculite, which appeared to create a different condition from that represented by phlogopite and muscovite. Understanding the strategy deployed by microorganisms to obtain mineral nutrients under different conditions provides new insights for interpreting the processes on which biodiversity and the homeostasis of soil environments depend.

## Supporting information

Supplementary Spreadsheet 1, Excel file containing the list of top up-regulated and down-regulated genes

Supplementary Spreadsheet 2, Excel file containing the list of GSEA results

## 6. Supplementary material

***Supplementary Methods:*** S1, MMN complete medium composition; S2, Confirmation of purity and identity of *P. involutus* strain; S3, Microcosm experiment; S4. Mineral substrates used for fungal weathering experiments; S5, SEM of mineral surfaces; S6, ICP-MS of mycelium; S7, RNA extraction; S8, Library preparation using TruSeq Stranded mRNA (Illumina) kit; S9, Bioinformatic analysis; S10, Statistical tests.

***Supplementary Tables:*** Supplementary Table S1, pH of agar under the mycelium; Supplementary Table S2, Average fungal biomass; Supplementary Table S3, Results of the sequencing of fungal mRNA.

***Supplementary Figures:*** Supplementary Figure S1, Diagram showing the setup of the experiment; Supplementary Figure S2, Volcano plots

***Supplementary Spreadsheets:*** Supplementary Spreadsheet 1, Excel file containing the list of top up-regulated and down-regulated genes; Supplementary Spreadsheet 2, Excel file containing the list of GSEA results.

## Availability of data and material (data transparency)

The datasets generated during and analysed during the current study are available in the Geo Submission Omnibus (GEO) public database with the accession number GSE158973.

## Acknowledgements

We thank Elena Lugli and Andie Hall for their daily support at the Molecular Laboratory, Emma Humphreys-Williams for her advice on chemical analyses, Luca Venturini and Matt Clark for helpful suggestions on bioinformatics solutions, Nathan Kenny for his suggestions on the manuscript, Innes Clatworthy and Alex Ball for support with SEM-EDS instrumentation.

## Authors’ contributions

F.P. conceived the design of the study, conducted molecular analyses, analysed and interpreted transcriptomic data and wrote the manuscript. A.J. conceived the design of the study, supervised the molecular and genetic part of project, and wrote the manuscript. J.C. conceived the design of the study, supervised the geochemical aspects of the project and wrote the manuscript.

## References

[1] Hoffland E, Kuyper TW, Wallander H, Plassard C, Gorbushina AA, Haselwandter K, et al. The role of fungi in weathering. Front Ecol Environ. 2004; 2: 258–264.

[2] Cuadros J. Clay minerals interaction with microorganisms: A review. Clay Miner. 2017; 52(2): 235–261.

[3] Fomina M, Skorochod I. Microbial Interaction with Clay Minerals and Its Environmental and Biotechnological Implications. Minerals. 2020; 10(10): 861.

[4] van der Heijden MGA, Martin FM, Selosse M-A, Sanders IR. Mycorrhizal ecology and evolution: the past, the present, and the future. New Phytol. 2015; 205: 1406–1423.

[5] Rosling A, Lindahl BD, Taylor AFS, Finlay RD. Mycelial growth and substrate acidification of ectomycorrhizal fungi in response to different minerals. FEMS Microbiol Ecol. 2004; 47(1): 31–37. https://doi.org/10.1016/S0168-6496(03)00222-8.

[6] Fomina M, Burford EP, Gadd G. Fungal dissolution and transformation of minerals: significance for nutrient and metal mobility. In: Gadd GM (ed.). Fungi in Biogeochemical Cycles. (Cambridge University Press, Cambridge) (ISBN: 978-0-521-84579-3). 2006. pp 236–266.

[7] Balogh-Brunstad Z, Kent Keller C, Thomas Dickinson J, Stevens F, Li C, Bormann BT. Biotite weathering and nutrient uptake by ectomycorrhizal fungus, *Suillus tomentosus*, in liquid-culture experiments. Geochim Cosmochim Acta. 2008; 72: 2601–2618.

[8] Bray AW, Oelkers EH, Bonneville S, Wolff-Boenisch D, Potts NJ, Fones GR, Benning LG. The effect of pH, grain size, and organic ligands on biotite weathering rates. Geochim Cosmochim Acta. 2015; 164: 127–145.

[9] Adeyemi AO, Gadd GM. Fungal degradation of calcium-, lead- and silicon-bearing minerals. Biometals. 2005; 18: 269–281.

[10] Cuadros J, Afsin B, Jadubansa P, Ardakani M, Ascaso C, Wierzchos J. Pathways of volcanic glass alteration in laboratory experiments through inorganic and microbially-mediated processes. Clay Miner. 2013; 48: 423–445.

[11] Pinzari F, Cuadros J, Napoli R, Canfora L, Baussà Bardají D. Routes of phlogopite weathering by three fungal strains. Fungal Biol. 2016; 120(12): 1582–1599.

[12] Schmalenberger A, Duran AL, Bray AW, Bridge J, Bonneville S, Benning LG, et al. Oxalate secretion by ectomycorrhizal *Paxillus involutus* is mineral-specific and controls calcium weathering from minerals. Sci Rep; 2015: 5: 1–14.

[13] Xiao B, Lian B, Sun L, Shao W. Gene transcription response to weathering of K-bearing minerals by *Aspergillus fumigatus*. Chem Geol. 2012; 306:1–9.

[14] Naranjo-Ortiz MA, Gabaldón T. Fungal evolution: major ecological adaptations and evolutionary transitions. Biol Rev Camb Philos Soc. 2019: 94(4): 1443–1476.

[15] Finlay RD, Mahmood S, Rosenstock N, Bolou-Bi EB, Köhler SJ, Fahad Z, et al. Reviews and syntheses: Biological weathering and its consequences at different spatial levels – from nanoscale to global scale. Biogeosciences. 2020; 17: 1507–1533.

[16] Hu A, Wang J, Sun H, Niu B, Si G, Wang J. et al. Mountain biodiversity and ecosystem functions: interplay between geology and contemporary environments. ISME J. 2020; 14: 931–944. https://doi.org/10.1038/s41396-019-0574-x

[17] Penman DE, Caves Rugenstein JK, Ibarra DE, Winnick MJ. Silicate weathering as a feedback and forcing in Earth’s climate and carbon cycle. Earth-Science Reviews. 2020; 209: 103298,

[18] Treseder KK, Lennon JT. Fungal traits that drive ecosystem dynamics on land. Microbiol Mol Biol Rev. 2015; 79(2): 243–262.

[19] Zanne A, Abarenkov K, Afkhami M, Aguilar-Trigueros CA, Bates S, Bhatnagar JM, et al. Fungal functional ecology: bringing a trait-based approach to plant-associated fungi. Biol Rev. 2020; 95(2): 409–433.

[20] Jargeat P, Chaumeton JP, Navaud O, Vizzini A, Gryta H. The *Paxillus involutus* (Boletales, Paxillaceae) complex in Europe: genetic diversity and morphological description of the new species *Paxillus cuprinus,* typification of *P. involutus* s.s., and synthesis of species boundaries. Fungal Biol. 2014; 118(1): 12–31. doi: 10.1016/j.funbio.2013.10.008.

[21] Kohler A, Kuo A, Nagy LG, Morin E, Barry KW, Buscot F, et al. Convergent losses of decay mechanisms and rapid turnover of symbiosis genes in mycorrhizal mutualists. Nat Genet. 2015; 47(4): 410–415. doi: 10.1038/ng.3223.

[22] Lapeyrie F, Chilvers GA, Bhem CA. Oxalic acid synthesis by the mycorrhizal fungus *Paxillus involutus* (Batsch EX FR) F.R. New Phytol. 1987; 106:139–146.

[23] Smits MM, Bonneville S, Benning LG, Banwart SA, Leake JR. Plant-driven weathering of apatite - the role of an ectomycorrhizal fungus. Geobiology. 2012; 10: 445–456.

[24] Saccone L, Gazze S, Duran A, Leake J, Banwart S, Ragnarsdottir K, et al., High resolution characterisation of ectomycorrhizal fungal-mineral interactions in axenic microcosm experiments. Biogeochemistry. 2012; 111: 411–425. doi:10.1007/s10533-011-9667-y.

[25] van Schöll L, Smits MM, Hoffland E. Ectomycorrhizal weathering of the soil minerals muscovite and hornblende. New Phytol. 2006; 171(4): 805–813.

[26] Wei Z, Kierans M, Gadd G. A Model Sheet Mineral System to Study Fungal Bioweathering of Mica. Geomicrobiol J. 2012; 29: 323–331.

[27] Dobin A, Davis CA, Schlesinger F, Drenkow J, Zaleski C, Jha S et al. STAR: ultrafast universal RNA-seq aligner. Bioinformatics. 2013; 29 (1): 15–21.

[28] Anders S, Pyl PT, Huber W, HTSeq—a Python framework to work with high-throughput sequencing data. Bioinformatics. 2015; 31(2): 166–169.

[29] Conesa A, Götz S, Garcia-Gomez JM, Terol J, Talon M, Robles M. Blast2GO: a universal tool for annotation, visualisation and analysis in functional genomics research. Bioinformatics. 2005; 21: 3674–3676.

[30] Conesa A, Götz S. Blast2GO: A Comprehensive Suite for Functional Analysis in Plant Genomics. Int J Plant Genomics. 2008; 1-13. 2008: 619832. doi: 10.1155/2008/619832.

[31] Robinson MD., McCarthy DJ. and Smyth GK. (20101). edgeR: a Bioconductor package for differential expression analysis of digital gene expression data. Bioinformatics (Oxford, England). 2010; 26(1): 139–140.

[32] Labarga A, Valentin F, Anderson M, Lopez R. Web Services at the European Bioinformatics Institute, Nucleic Acids Res. 2007; 35 (Issue suppl-2): W6–W11. https://doi.org/10.1093/nar/gkm291.

[33] Huerta-Cepas J, Forslund S, Coelho LP, Szklarczyk D, Jensen L, von Mering C, Bork P. Fast Genome-Wide Functional Annotation through Orthology Assignment by eggNOG-Mapper. Mol Biol Evol. 2017; 34. 10.1093/molbev/msx148.

[34] Subramanian A, Tamayo P, Mootha VK, Mukherjee S, Ebert BL, Gillette MA, et al. Gene set enrichment analysis: A knowledge-based approach for interpreting genome-wide expression profiles. PNAS. 2005; 102(43): 15545–15550.

[35] Chen Y, Meltzer PS. Gene Expression Analysis via Multidimensional Scaling. Current Protocols in Bioinformatics. 2005; Chapter 7: Unit 7.11. doi: 10.1002/0471250953.bi0711s10.

[36] Rodriguez-Navarro, A. Potassium transport in fungi and plants. Biochim Biophys Acta. 1999; 1469 (2000): 1–30.

[37] Camacho M, Ramos J, Rodríguez-Navarro A. Potassium requirements of *Saccharomyces cerevisiae*. Curr Microbiol. 1981; 6: 295–299.

[38] Gostinčar C, Lenassi M, Gunde-Cimerman N, Plemenitaš A. Fungal adaptation to extremely high salt concentrations. Adv Appl Microbiol. 2011; 77: 71–96.

[39] Garcia K, Delteil A, Conejero G, Becquer A, Plassard C, Sentenac H, Zimmermann S. Potassium nutrition of ectomycorrhizal *Pinus pinaster*: overexpression of the *Hebeloma cylindrosporum* HcTrk1 transporter affects the translocation of both K and P in the host plant. New Phytol. 2014; 201: 951–960.

[40] Jentschke G, Brandes B, Kuhn AJ, Schröder WH, Godbold DL. Interdependence of phosphorus, nitrogen, potassium and magnesium translocation by the ectomycorrhizal fungus *Paxillus involutus*. New Phytol. 2001; 149: 327–337.

[41] Bücking H, Heyser W. Elemental composition and function of polyphosphates in ectomycorrhizal fungi — an X-ray microanalytical study. Mycol Res. 1999; 103(1): 31–39.

[42] Orlovich DA, Ashford AE. Polyphosphate granules are an artefact of specimen preparation in the ectomycorrhizal fungus *Pisolithus tinctorius*. Protoplasma. 1993; 173: 91–105.

[43] Ashford AE, Vesk PA, Orlovich DA, Markovina A-L, Allaway WG. Dispersed polyphosphate in fungal vacuoles of *Eucalyptus pilularis/Pisolithus tinctorius* ectomycorrhizas. Fungal Genet Biol. 1999; 28: 21–33.

[44] Bonneville SC, Bray A, Benning L. Structural Fe(II) Oxidation in Biotite by an Ectomycorrhizal Fungi Drives Mechanical Forcing. Environ sci & technol. 2016; 50(11): 5589–5596.

[45] Väre H. Aluminium polyphosphate in the ectomycorrhizal fungus *Suillus variegatus* (Fr.) O. Kunze as revealed by energy dispersive spectrometry. New Phytol. 1990; 116: 663–668.

[46] Gadd GM. Metals, minerals and microbes: geomicrobiology and bioremediation. Microbiology. 2010; 156(3): 609–643.

[47] Illmer P, Buttinger R. Interactions between iron availability, aluminium toxicity and fungal siderophores. Biometals. 2006; 19(4): 367–377.

[48] Haro R, Benito B. The Role of Soil Fungi in K+ Plant Nutrition. Int J Mol Sci. 2019; 20(13): 3169. doi:10.3390/ijms20133169.

[49] Dueñas-Santero E, Martín-Cuadrado AB, Fontaine T, Latgé J P, del Rey F, Vázquez de Aldana C. Characterization of glycoside hydrolase family 5 proteins in *Schizosaccharomyces pombe*. Eukaryotic cell. 2010; 9(11): 1650–1660.

[50] Paparokidou C, Leake JR, Beerling DJ et al. Phosphate availability and ectomycorrhizal symbiosis with *Pinus sylvestris* have independent effects on the *Paxillus involutus* transcriptome. Mycorrhiza. 2012; 31: 69–83.

[51] Lian B, Wang B, Pan M, Liu C, Teng HH. Microbial release of potassium from K-bearing minerals by thermophilic fungus *Aspergillus fumigatus*. Geochim Cosmochim Acta. 2008; 72:87–98.

[52] Bonneville S, Smits MM, Brown A, Harrington J, Leake JR, Brydson R, Benning LG. Plant-driven fungal weathering: Early stages of mineral alteration at the nanometer scale. Geology. 2009; 37(7): 615–618.

[53] Ahmed E, Holmström SJM. Microbe-mineral interactions: The impact of surface attachment on mineral weathering and element selectivity by microorganisms. Chem Geol. 2015; 403: 13–23.

[54] van Hees PA, Rosling A, Essén S, Godbold DL, Jones DL, Finlay RD. Oxalate and ferricrocin exudation by the extramatrical mycelium of an ectomycorrhizal fungus in symbiosis with *Pinus sylvestris*. New Phytol. 2006; 169(2): 367–77. doi: 10.1111/j.1469-8137.2005.01600.x.

[55] Martínez AT, Ruiz-Dueñas FJ, Camarero S, Serrano A, Linde D, Lund H et al. Oxidoreductases on their way to industrial biotransformations. Biotechnol Adv. 2017; 35(6); 815–831.

[56] Song M, Pedruzzi I, Peng Y, Li P, Liu J-F, Yu J. K-Extraction from Muscovite by the Isolated Fungi. Geomicrobiol J. 2015; 32(9): 771–779.

[57] Xiao L, Sun Q, Lian B. A Global View of Gene Expression of *Aspergillus nidulans* on Responding to the Deficiency in Soluble Potassium. Curr Microbiol. 2016; 72(4): 410–419.

[58] Wang W, Lian B, Pan L. An RNA-sequencing study of the genes and metabolic pathways involved in *Aspergillus niger* weathering of potassium feldspar. Geomicrobiol J. 2015; 32(8): 689–700.

[59] Wang W, Sun Q, Lian B. Redox of Fungal Multicopper Oxidase: A Potential Driving Factor for the Silicate Mineral Weathering, Geomicrobiol J. 2018; 35(10): 879–886.

[60] Garcia K and Zimmermann SD. The role of mycorrhizal associations in plant potassium nutrition. Front. Plant Sci. 2014; 5: 337. doi: 10.3389/fpls.2014.00337

[61] Benito B, Garciadeblás B, Fraile-Escanciano A et al. Potassium and sodium uptake systems in fungi. The transporter diversity of *Magnaporthe oryzae*. Fungal Genet Biol. 2011; 48: 812–822.

[62] Corratgé C, Zimmermann S, Lambilliotte R, Plassard C, Marmeisse R, Thibaud JB, et al. Molecular and functional characterization of a Na+-K+ transporter from the Trk family in the ectomycorrhizal fungus *Hebeloma cylindrosporum*. J. Biol. Chem. 2007; 282: 26057–26066.

[63] Heaton LLM, Jones NS, Fricker MD. A mechanistic explanation of the transition to simple multicellularity in fungi. Nat Commun. 2020; 11: 2594. https://doi.org/10.1038/s41467-020-16072-4.

