## Supplementary Methods, Tables and Figures for "Fungal taste for minerals: the ectomycorrhizal fungus *Paxillus involutus* triggers specific genes when extracting potassium from different silicates"

#### *S1. MMN complete medium composition*

Modified Melin– Norkrans (MMN) medium (composition: 10g $l^{-1}$ glucose, 250 mg l−1 $KH_2PO_4$, 250 mg l−1 $(NH_4)2HPO_4$, 150 mg l−1 $MgSO_4 \times 7H_2O$, 25 mg l−1 NaCl, 50 mg l−1 $CaCl_2 \times 2H_2O$, 12 mg l−1 $FeCl_3 \times 6H_2O$ and 1 mg l−1 thiamine-HCl; trace elements, pH 5.6) modified from Langer et al., (2008).

#### *S2. Confirmation of purity and identity of P. involutus strain*

Before undertaking the experiments, the purity and identification of the fungal strain reactivated from freeze-dried ATCC culture was verified by DNA extraction and sequencing of ITS1 and ITS2 regions, and the adjacent 5.8S. DNA extraction was performed directly from fungal mycelium using the PowerLyzer PowerSoil DNA isolation Kit (catalogue n. 12855, UK, QIAGEN Ltd.). The extracted DNA was visualised running electrophoresis on a 1% agarose gel and used for PCR amplification. Fragments of 450-600 bp in size were amplified with the primer pair ITS1 and ITS4, according to White et al. (1990). PCR reactions were performed with the BioTAQ® polymerase (Bioline®). A mixture was prepared with 50 units $ml^{-1}$ of Taq DNA Polymerase supplied in 10xNH4-based reaction buffer at pH 8.5, 400 µM dATP, 400 µM dGTP, 400 µM dCTP, 400 µM dTTP, 3 mM MgCl2, and diluted to 1 x before its use. 12.5 pmol $\mu l^{-1}$ of each primer (stock: 50 pmol $\mu l^{-1}$, Sigma Aldrich, UK) was added to the mix. PCR was performed in an Applied Biosystems® Veriti® 96-Well Thermal Cycle. The thermocycling program was 3 min denaturation at 94° C, followed by 35 cycles of 30 sec. denaturation at 94° C, 30 sec. annealing at 55° C, and 1 min extension at 72° C. Ten minutes at 72° C were used as a final extension step. PCR products were purified using the GenElute™ PCR Clean-Up Kit (catalogue n.NA1020, Sigma Aldrich, UK), following the manufacturer protocol and eluting the purified DNA in 50 µl of ddH2O water. The purified PCR products were Sanger sequenced at the NHM Sequencing Facility. The forward and reverse electropherograms obtained were manually edited and aligned using the BioEdit software program (http://www.mbio.ncsu.edu/BioEdit/bioedit.html) to obtain a consensus sequence that was compared using the BLAST search program with NCBI (Altschul et al., 1990) and UNITE databases (Nilsson et al., 2018).

#### *S3. Microcosm experiment*

The microcosms were prepared using round sterile polystyrene 100x20 mm Petri dishes (N. 430167 Corning). 30 ml of agar were poured in each Petri dish. The positive control consisted of microcosms made with full MMN modified medium while the negative control and the microcosms containing the minerals were prepared using K-depleted MMN agar, prepared without $KH_2PO_4$. The purchased reagents to prepare K-depleted Agar medium contained traces of K, since a concentration of 0.3 mg $kg^{-1}$ K in final K-depleted Agar was measured. 20 biological replicates (microcosms) were prepared for each experiment. Sterile cellophane membranes (Cellophane™ 325P, 80mmm DIA, SBC292529

AA Packaging Ltd, Lancashire, UK) were placed on top of the agar before adding fungi and minerals. The membrane was essential to harvest the fungal biomass without residual agar. Mineral flakes were placed in the centre of the Petri dish, on the membrane, and the fungus was inoculated in three points, around them, in order to ensure contact between the minerals and growing mycelium (Wei et al., 2012). The inoculums consisted of 5 mm in diameter agar plugs of *P. involutus*, obtained from the edge of 7-day-old actively growing cultures. All cultures were kept in the dark at 25˚C throughout the experiment. The microcosms were incubated for 21 days. Ten biological replicates from each experiment were used for RNA extraction, five from each experiment were dismounted and used for inductively coupled plasma (ICP) analysis, and the corresponding minerals were used in scanning electron microscopy (SEM) analysis. After the experiments, fungal biomass was determined by weighing the whole mycelium from 5 microcosms from each experiment and weighing the biomass again after oven-drying (50°C, 48h). The pH of the agar was measured at the beginning and the end of the experiments (WWR Phenomenal pH-meter with electrode n.662-1164).

***S4. Mineral substrates used for fungal weathering experiments***

The minerals used, with decreasing level of K weatherability, and their structural formulas are (SEM-EDX and microprobe analyses): muscovite $(Si_{3.13}Al_{0.87})_{tet}$ $(Al_{1.72}Mg_{0.11}Fe^{3+}{}_{0.19})_{oct}$ $(Na_{0.10}K_{0.83})_{int}$ $O_{10}$ $(OH)_2$; phlogopite: $(Si_{2.75}Al_{1.25})_{tet}$ $(Al_{0.14}Mg_{2.09}Fe^{2+}{}_{0.49}Ti_{0.17})_{oct}$ $(K_{0.98})_{int}$ $O_{10}$ $(OH)_2$; and K-vermiculite: $(Si_{2.96}Al_{0.81}Fe^{3+}{}_{0.23})_{tet}$ $(Mg_{2.86}Fe^{3+}{}_{0.09}Ti_{0.05}Cr_{0.01})_{oct}$ $(Mg_{0.23}Ca_{0.01}K_{0.38})_{int}$ $O_{10}$ $(OH)_2$, where tet, oct and int stand for tetrahedral, octahedral and interlayer sites in the crystals. X-ray-diffraction did not detect other associated minerals. The minerals were cut as flakes of ~2 $cm^2$, cleaned ultrasonically in deionised water, and sterilised by autoclaving (121°C, 15 minutes). The K-vermiculite was prepared by cation exchange in concentrated KCl solution (4.42 g/100 ml, 14 days under agitation).

***S5. SEM of mineral surfaces***

Mineral samples were analysed before and after the experiment using two microscopes. Uncoated samples were studied with a variable pressure SEM instrument (EVO50, Carl-Zeiss Electron Microscopy Group); and Au-Pd-coated samples were studied in a Zeiss Ultra Plus Field Emission SEM for high-resolution imaging. Voltage was always 20 kV.

***S6. ICP-MS of mycelium***

Groups of five replicates from each experiment were used to analyse Na, Mg, Al, Fe, P, K, Ca, Fe in the fungal mycelium. The fungus was peeled off the agar plates together with the supporting membrane, paying attention not to carry along agar fragments. The minerals were removed from the agar in the treatments. The fungal biomass was weighed on a precision scale (four decimal places) and transferred to a 50 mL Polypropylene tube. Samples were digested and analysed by inductively coupled plasma mass spectrometry (ICP-MS, Agilent Technologies 7700). Digestion was initiated by adding 2.0 mL $HNO_3$, and 0.5mL $H_2O_2$ 30%. Caps were hand-tightened and tubes were vortexed to ensure the entire sample was wetted and placed into the digestion block and heated at 70°C, 48 h, degassing after the first 2 hours. Pressure build-up in the tubes during the initial 30 min warming period was released by loosening each cap sufficiently to equalise pressure, then immediately re-tightening firmly and replacing on the digestion block. Digested samples were removed from the digestion block and cooled to room temperature. Samples were made to 10 ml volume with $ddH_2O$. No undissolved material was observed.

***S7. RNA extraction***

At the end of the experiment, fungal biomass was removed from the cellophane membrane and ground into a powder in liquid nitrogen using a sterilised mortar and pestle. Up to 200 mg of frozen biomass from each biological replicate was extracted using the RNAeasy® PlantMini Kit (Qiagen) using RLC buffer and on-column DNAse treatment according to the manufacturer's instructions. The absence of genomic DNA in the extracts was confirmed by PCR amplification of ITS1 and ITS2 regions with the primer pair ITS1 and ITS4 according to White et al. (1990), as described in supplementary paragraph 2 and visualised on 1% agarose gels by electrophoresis.

Total RNA was eluted in 50 µl of sterile, distilled and RNAse free $H_2O$ and stored at -80°C. The concentrations and quality of purified nucleic acids were checked on a NanoDrop (ND-8000 NanoDrop Technologies), Qubit® (using the RNA HS Assay Kits), and Agilent TapeStation 2200 using the High Sensitivity RNA ScreenTape (Agilent).

***S8. Library preparation using TruSeq Stranded mRNA (Illumina) kit***

For each experiment, the RNA extracts from five biological replicates (microcosms) were sequenced. RNA-seq transcriptome libraries were prepared with a TruSeq Stranded mRNA Kit following the manufacturer's instructions (Illumina, TruSeq Stranded mRNA Reference Guide, Document # 1000000040498 v00, October 2017) from 1 µg of total RNA. Poly-A containing mRNA molecules were purified using the Dynabeads® mRNA Purification Kit containing poly-T oligo attached magnetic beads (Ambion, Life Technologies). The double-stranded cDNA was then synthesised from 100 ng mRNA. Cleaved RNA fragments were copied into first-strand cDNA using reverse transcriptase and random primers. Actinomycin D was added to FSA (First Stand Synthesis Act D mix) to prevent spurious DNA-dependent synthesis, while allowing RNA-dependent synthesis, improving strand specificity. Strand specificity was achieved by replacing dTTP with dUTP in the SMM (Second Strand Marking Mix), followed by second-strand cDNA synthesis using DNA Polymerase I and RNase H. The synthesised cDNA was subjected to end-repair, phosphorylation and 'A' base addition, according to Illumina library construction protocol. A total of 25 libraries (5 biological replicates for each experiment) were sequenced with Illumina NovaSeq 6000 (2 x 100 bp read-length) by the CeGAT Company (Germany).

***S9. Bioinformatic analysis***

Demultiplexing of the sequencing reads was performed with Illumina bcl2fastq (version 2.19). Adapters were trimmed with Skewer (version 0.2.2) (Jiang et al. 2014). The quality of FASTQ files was analysed with FastQC (version 0.11.5-cegat) (Andrews 2010). Trimmed reads were aligned to the *Paxillus involutus* reference genome (GCA_000827475.1) using STAR (v2.5.2b, Dobin et al. 2013), and saved as BAM files (one per sample). A count table was created using the HTSeq package (Anders et al., 2014) implemented in OmicBox (version 1.2.4). Reads were only considered when they mapped unambiguously to a single genomic feature, and multimapping reads or those overlapping with more than one feature were discarded.

The count table was used to perform pairwise differential expression analysis (DEA) between pairs of experimental conditions (with 5 replicates per experimental condition). EdgeR version 2.12 (Robinson et al., 2010) implemented in OmicsBox (version 1.2.4) was used in this step. The minimum CPM (count-per-million) filter was set at 2, with a minimum number of samples reaching this CPM Filter set at 3 (to exclude genes with low counts across libraries, performed on a CPM basis to account for differences in library size). The normalisation method used was TMM (Trimmed mean of M values: weights are obtained from the delta method on Binomial Data), and the selected statistical test was the Exact Test (robust). "log2-fold-changes" (logFC) were used to measure extent of expression changes between conditions. These results were corrected for False Discovery Rate (FDR) between conditions using the Benjamini-Hochberg method (multiple hypothesis testing corrections). A gene was considered upregulated or downregulated when FDR-corrected p-value $\leq 0.05$, and logFC $> 1$ or downregulated when FDR-corrected p-value $\leq 0.05$, and logFC $< -1$. Volcano plots of differentially expressed genes (DEG) were generated using OmicsBox (version 1.2.4).

A heatmap was built using ComplexHeatmap package implemented in Bioconductor (Gu et al., 2016). The expression data were clustered according to Ward's method, complete-link clustering.

Rows were scaled according to the standard procedure proposed in Bioconductor (Hahne et al., 2010). After scaling, all values were transformed so that negative values were related to downregulation and positive values to upregulation.

Multidimensional scaling (MDS) was used to visualise the level of similarity between the experiments and the biological replicates. The percentage values on the axes describe how much of the variance between samples is captured in a Cartesian space (Chen and Meltzer 2005).

***S10. Statistical tests***

One-way analysis of the variance (ANOVA) was applied when comparing the concentration of elements, fungal biomass and pH in the samples, and the significance of the differences was tested at 99% confidence. ANOVA was followed by a post-hoc analysis using Tukey's test. The Pearson correlation coefficient was used as measure of linear correlation between elements' concentration in the samples. Pearsons's "r" has a value between +1 and −1. A value of +1 is total positive linear correlation, 0 is no linear correlation, and −1 is total negative linear correlation. Significance of the correlation was tested using a permutation test (XLSTAT statistical software Version 2019.3.2, Addinsoft 1995-2020, France).

**Supplementary Tables**

**Supplementary Table S1.** pH of agar under the mycelium at the end of 21 days experiment. Values are the means from n = 5 replicates. Different letters indicates in each column statistically significant differences (P < 0.05), one-way ANOVA

| **(initial agar pH 4.7**±0.05) | pH |
|---|---|
| Cp | 3.00±0.36 a |
| Cm | 2.54±0.42 ab |
| M | 2.12±0.19 b |
| P | 2.12±0.09 b |
| V | 2.27±0.33 b |
| Pr > F(Model) | 0.0022 |
| Significant | Yes |

**Supplementary Table S2.** Average fungal biomass from each experiment after 21 days incubation. Values are the means from n = 5 replicates. Different letters indicate in each column statistically significant differences (P < 0.05), one-way ANOVA.

| | **Wet weight (g)** | **Dry weight (g)** | **Water content (g)** |
|---|---|---|---|
| Cp | 0.67±0.03 a | 0.11±0.01 ab | 0.57±0.03 a |
| Cm | 0.43±0.03 c | 0.10±0.01 b | 0.33±0.02 c |
| M | 0.44±0.04 c | 0.10±0.01 ab | 0.34±0.03 c |
| P | 0.54±0.04 b | 0.11±0.01 a | 0.43±0.03 b |
| V | 0.48±0.03 c | 0.11±0.01 ab | 0.37±0.02 c |
| Pr > F(Model) | < 0.0001 | 0.03 | < 0.0001 |
| Significant | Yes | Yes | Yes |

**Supplementary Table S3.** Results of the sequencing of fungal mRNA. Five biological replicates were used in each experiment: Cm, positive control; Cp, negative control; M, muscovite; P, phlogopite; V, vermiculite.

| | **Mean raw read pairs per replicate (million)** | | | **Mean number of bases (in Gbp)** | | |
|---|---|---|---|---|---|---|
| | **Mean** | | | **Mean** | | |
| **Cm** | 37.89 | ± | 2.67 | 38.18 | ± | 7.64 |
| **Cp** | 36.13 | ± | 6.18 | 36.40 | ± | 7.28 |
| **M** | 39.01 | ± | 2.53 | 39.27 | ± | 7.85 |
| **P** | 38.88 | ± | 4.18 | 39.20 | ± | 7.84 |
| **V** | 41.48 | ± | 6.05 | 41.83 | ± | 8.37 |

### Supplementary Figures

**Supplementary Figure S1.** Diagram showing the setup of the experiment. The microcosms were prepared using round sterile polystyrene Petri dishes. The positive control (Cp) consisted of microcosms made with full MMN modified medium while the negative control (Cm) and the microcosms containing the minerals were prepared using K-depleted MMN agar. Sterile cellophane membranes were placed on top of the agar before adding fungi and minerals. Mineral flakes were placed in the centre of the Petri dish, on the membrane, and the fungus was inoculated in three points, around them.

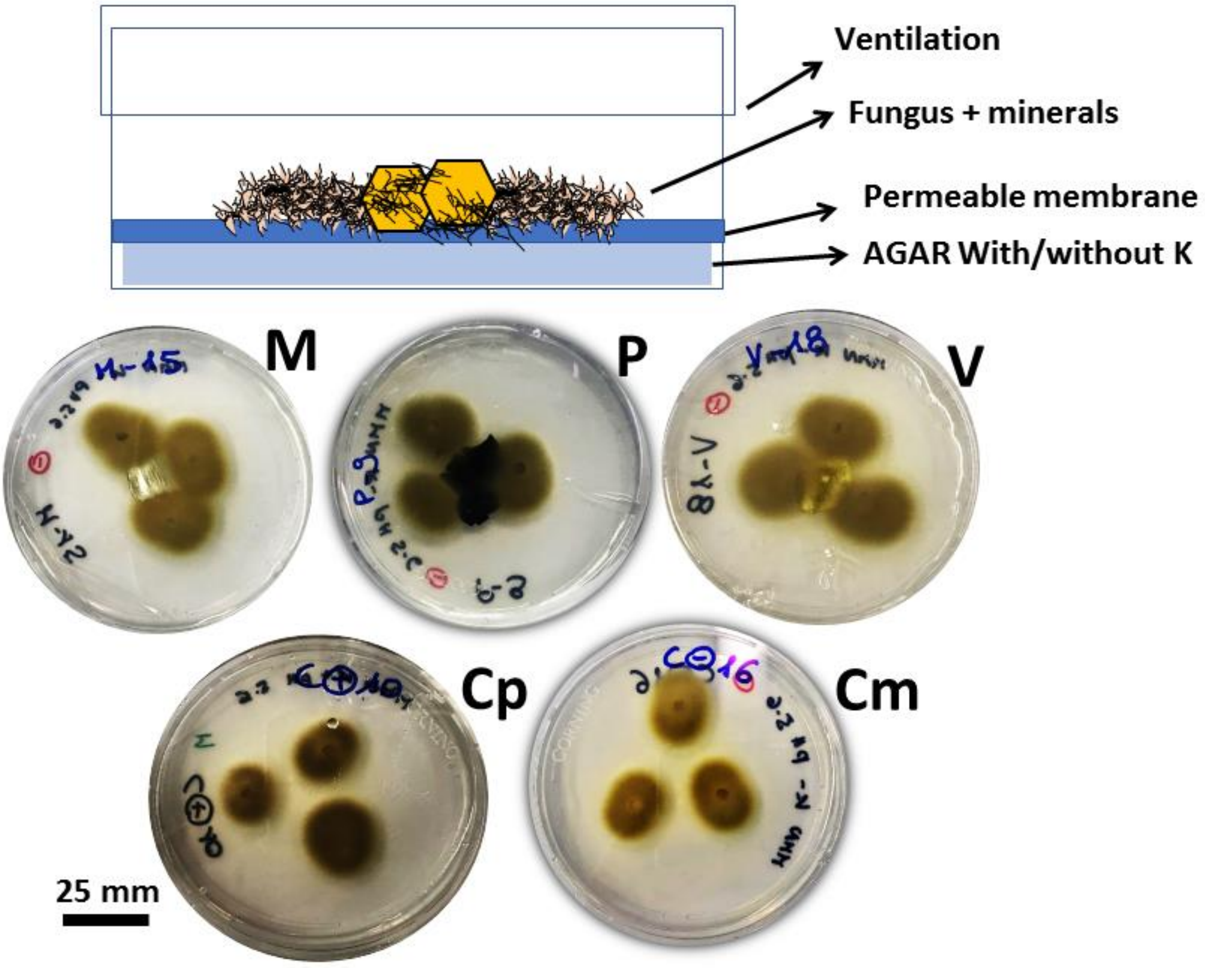

**Supplementary Figure S2.** Volcano plots of genes differentially expressed between selected treatments. The differently expressed genes are visualised as dots in a system where the X-axis represents the log2 of the fold changes, and the Y-axis the -log10 (p-value) of the significance: the green dots indicate up-regulated DEGs; the red dots represent down-regulated DEGs; the black dots represent genes that are not differentially expressed between treatments; the grey dots are differentially expressed but not up or down regulated at the chosen p-value. Scatter chart is constructed by plotting the negative log of the adjusted p-values (FDR) on the y-axis versus the log of the fold changes on the x-axis.

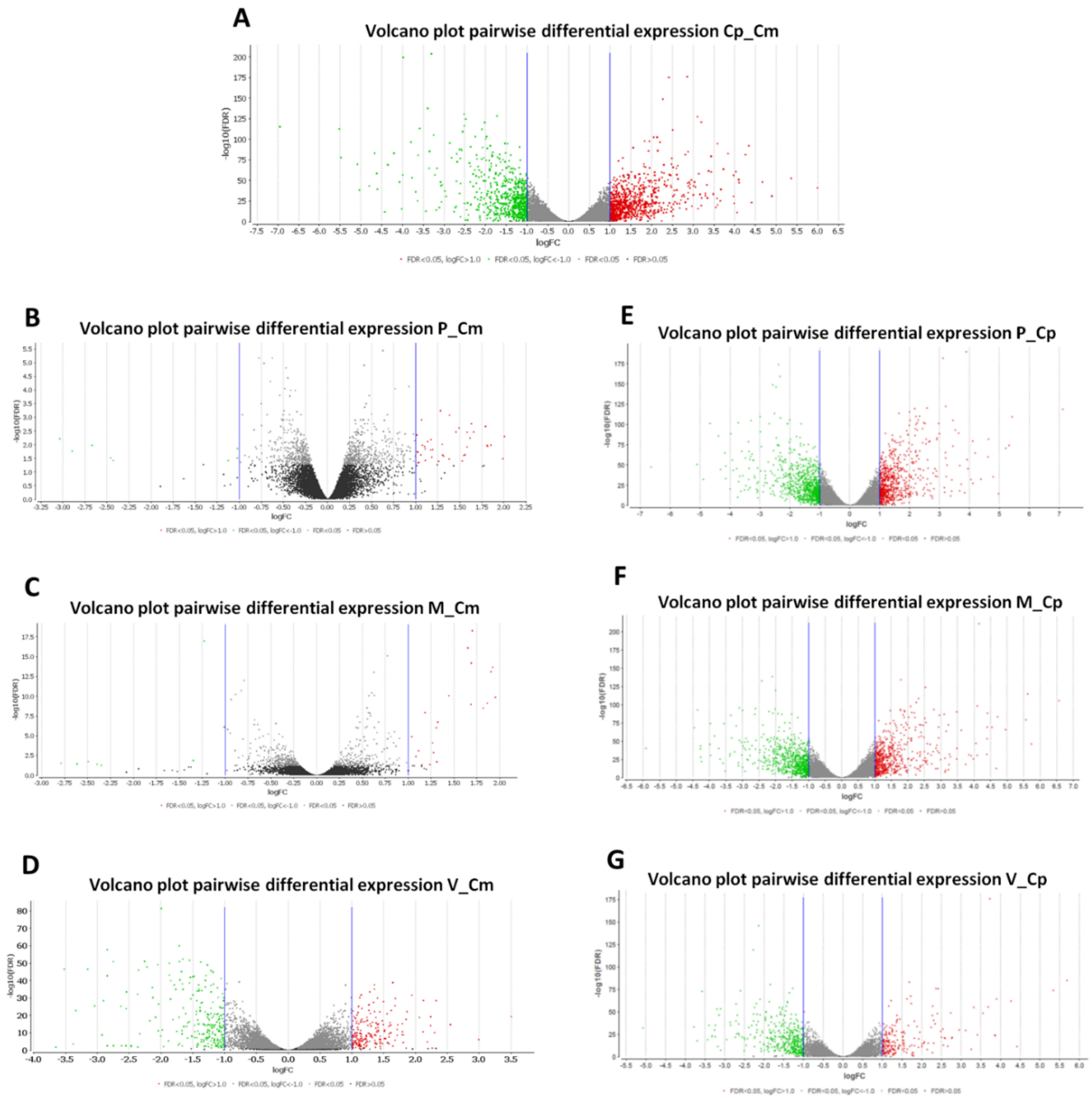

**Supplementary Spreadsheets**

**Supplementary Spreadsheet 1:** Excel file containing the list of top up-regulated and down-regulated genes between the contrasts: Cp/Cm, V/Cm, P/Cm and M/Cm. In the file the following information are reported: type of contrast, *Paxillus involutus* gene name, logFC values, FDR values, gene function description, gene length, GO IDs, process GO names, function GO names and cellular component GO names.

**Supplementary Spreadsheet 2:** Excel file containing the list of GSEA results between the contrasts: Cp/Cm, V/Cm, P/Cm and M/Cm. In each table the following information are reported: Tags (top or bottom), GO ID, GO Name, GO Category, size (number of genes in the enriched category), ES values, NES values, nominal p-value, FDR values, q-val, FWER (Family-Wise Error Rate), p-value, rank at max, leading edge, core enrichment sequences, no-core enrichment sequences (Subramanian et al., 2005).

***References***


Altschul SF, Gish W, Miller W, Myers EW, Lipman DJ. Basic local alignment search tool. J Mol Biol. 1990; 5: 215(3): 403–410.

Anders S, Pyl PT, Huber W. HTSeq — A Python framework to work with high-throughput sequencing data. Bioinformatics. 2014; 31(2): 166–169.

Andrews S. FastQC A Quality Control tool for High Throughput Sequence Data. 2010; http://www.bioinformatics.babraham.ac.uk/projects/fastqc

Chen Y, Meltzer PS. Gene Expression Analysis via Multidimensional Scaling. Current Protocols in Bioinformatics. 2005; 10: 7.11.1–7.11.9.

Gu Z, Eils R, Schlesner M. Complex heatmaps reveal patterns and correlations in multidimensional genomic data. Bioinformatics. 2016; 15; 32(18): 2847–2849.

Hahne F, Huber W, Gentleman R, Falcon S. Bioconductor Case Studies. Use R! series. Springer Science & Business Media, New York, 2010, pp.284.

Jiang H, Lei R, Ding SW, Zhu S. Skewer: a fast and accurate adapter trimmer for next-generation sequencing paired-end reads. BMC Bioinformatics. 2014; 12: 15: 182. doi: 10.1186/1471-2105-15-182.

Langer D, Krpata U, Peintner W, Wenzel W, Schweiger P. Media formulation influences in vitro ectomycorrhizal synthesis on the European aspen *Populus tremula* L. Mycorrhiza. 2008; 18(6-7): 297–307.

Nilsson RH, Larsson K-H, Taylor AFS, Bengtsson-Palme J, Jeppesen TS, Schigel D, *et al.* The UNITE database for molecular identification of fungi: handling dark taxa and parallel taxonomic classifications. Nucleic Acids Res. 2018; 47(D1): D259–D264.

Robinson MD, McCarthy DJ, Smyth GK. edgeR: a Bioconductor package for differential expression analysis of digital gene expression data. Bioinformatics (Oxford, England). 2010; 26(1): 139–140.

Subramanian A, Tamayo P, Mootha VK, Mukherjee S, Ebert,BL, Gillette MA, *et al.* Gene set enrichment analysis: A knowledge-based approach for interpreting genome-wide expression profiles. PNAS. 2005; 102(43): 15545–15550.

Wei Z, Kierans M, Gadd G. A Model Sheet Mineral System to Study Fungal Bioweathering of Mica. Geomicrobiol J. 2012; 29: 323–331.

White TJ, Bruns T, Lee S, Taylor J. Amplification and direct sequencing of fungal ribosomal RNA genes for phylogenetics. In: Innis M, Gelfand D, Sninsky J, White TJ (eds.). PCR Protocols: A Guide to Methods and Applications. Chapter 38. New York: Academic Press, 1990, pp. 315–322.
